# Autophagy-dependent proteome remodelling and ribosome decline balance starvation survival and recovery speed in *C. elegans*

**DOI:** 10.1101/2024.08.28.609383

**Authors:** Joel Tuomaala, Devanarayanan Siva Sankar, Julie Perey, Sacha Psalmon, Klement Stojanovski, Nicholas Stroustrup, Joern Dengjel, Benjamin D. Towbin

## Abstract

Animals facing fluctuating food availability must balance survival during starvation with rapid resumption of growth when encountering food. We investigated how proteome turnover and remodelling through autophagy influences this trade-off in *C. elegans* L1 larvae by combining live imaging and proteomics. Starvation triggered an autophagy-dependent, disproportionate loss of ribosomal and other growth-related proteins. Residual ribosomal protein abundance at the end of starvation predicted the rate of growth recovery of individual animals during post-starvation feeding, linking proteome-scale changes to organism-scale life-history. Hyperactivation of the mTORC1 regulator RAGA-1 preserved ribosomal proteins, accelerated recovery after short starvation, but reduced survival under prolonged starvation. These findings identify autophagy-dependent ribosomal protein decline as a central component of starvation-induced proteome remodelling and reveal its role in balancing the trade-off between starvation endurance and recovery speed in a multicellular animal.

## Introduction

Nutrient fluctuations impose a dual challenge for animals: enduring prolonged starvation and resuming growth efficiently when food returns. Balancing these tasks is essential for fitness in environments where food availability is unpredictable ^1–3^.

During starvation, eukaryotes shut down most protein synthesis and recycle existing proteins via autophagy ^4,5^ or proteasomal degradation ^6,7^. This proteome recycling supplies energy and metabolites but can also remove components essential for survival ^8^. One way to limit such costs is to preferentially degrade proteins that are dispensable during starvation. Selective autophagy can target specific structures, such as the endoplasmic reticulum or the Golgi apparatus, via dedicated receptor proteins ^9^. Ribosome, too, have been observed to decline under nutrient stress and recent work has begun to uncover the pathways mediating this ribosomal turnover ^10–16^. Yet, to what extent such non-uniform turnover of the proteome at the scale of an entire animal shapes the tradeoff between starvation survival and growth is unknown. We addressed this question using the *C. elegans* L1 diapause, a developmental stage that naturally withstands prolonged starvation ^4^ and is critical for fitness during the boom-bust cycles frequently experienced by *C. elegans* in the wild ^17^.

We combined quantitative proteomics with live imaging to track proteome composition and growth recovery dynamics in individual animals. We observed a systematic autophagy-dependent decline and remodelling of the proteome during starvation: cytoplasmic, mitochondrial, and endoplasmic reticulum proteins declined in relative abundance, with ribosomes and other translation factors showing the most pronounced reduction. Ribosomal protein abundance at the end of starvation correlated with recovery speed, suggesting that ribosomes act as a limiting factor for post-starvation growth. Manipulating nutrient-sensing pathways confirmed a functional relation between proteome change and recovery speed. Hyperactivation of the mTORC1 activator RAGA-1 slowed proteome remodelling, reduced ribosomal protein loss, accelerated recovery after short starvation, and reduced survival after prolonged starvation. These results demonstrate a progressive shift of the proteome in its composition during starvation and suggest that this shift impacts the survival and recovery trade-off.

## Results

### Growth-related proteins decline rapidly during starvation

To measure proteome dynamics during starvation, we conducted quantitative label-free proteomics ^18^ of L1 larvae of *C. elegans* after 1 and 9 days of starvation (Fig. 1a). Using algorithms optimized for within (iBAQ ^19^) and across (LFQ ^20^) sample comparisons, we detected 2624 and 1787 proteins, respectively, across time points and repeats. Ribosomal proteins accounted for 12.1% ± 0.2% (s.e.m.) of the total detected protein mass on day 1 and declined to 9.0% ± 0.3%, i.e. by 27%, by day 9. The total detected mitochondrial protein mass declined to a similar degree (-23.8%), non-ribosomal cytoplasmic proteins, however, declined less strongly (12.5%), whereas nuclear, filamentous and membrane bound proteins increased in relative abundance (Fig. 1d). Since these measurements represent relative changes in the fractional proteome composition, the decline in ribosomal proteins is not explained by overall body shrinkage, but by a relative difference in production or decay compared to other proteins. Similarly, an increase in relative protein abundance, as observed for nuclear proteins, can result from comparatively slower degradation or from *de novo* synthesis.

**Figure 1.**
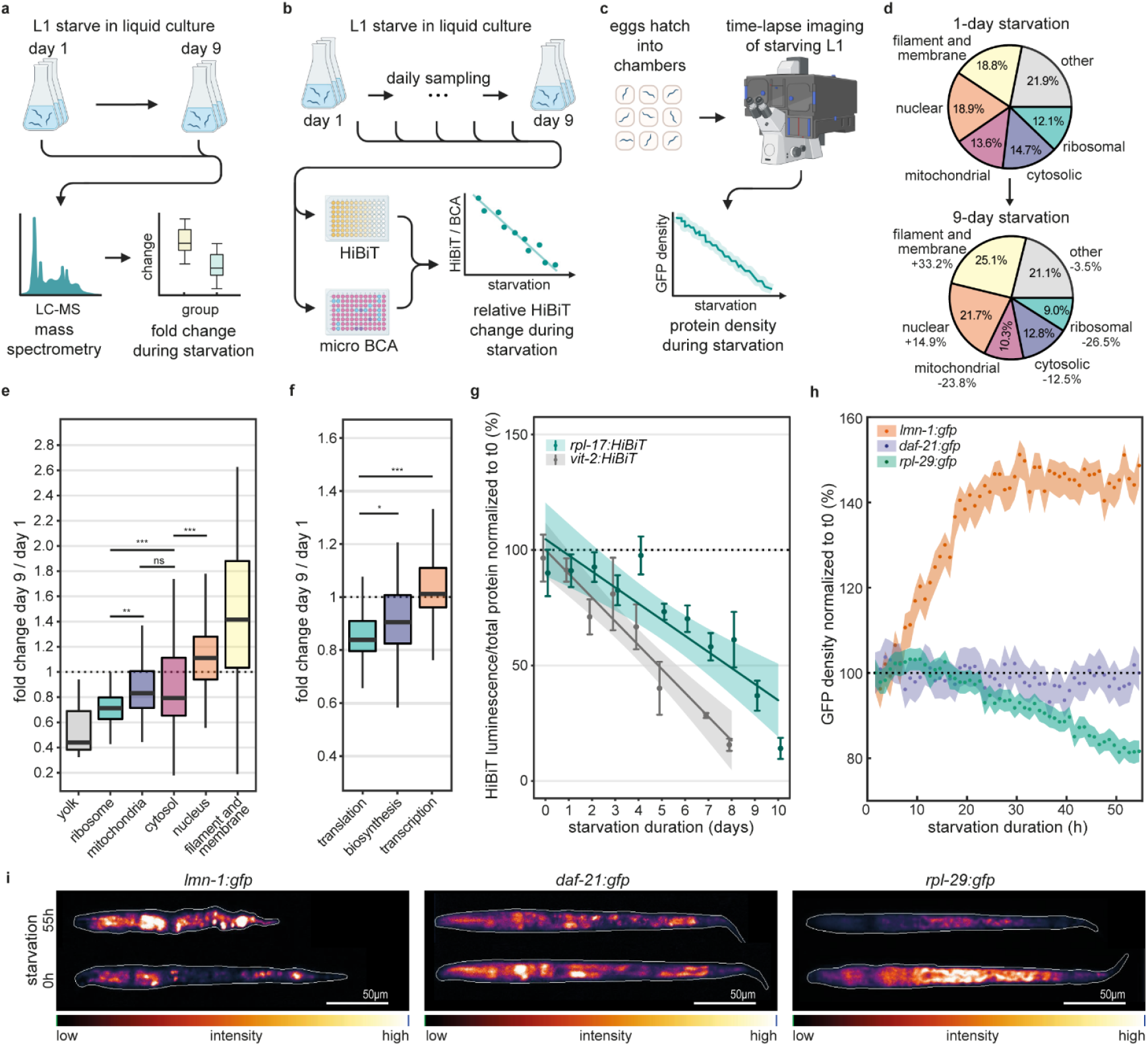
Rapid ribosomal protein decline during starvation. **a-c**. Experimental design of measurements of protein dynamics during starvation of L1 larvae by mass spectrometry (a) HiBiT (b) and live imaging (c). **d**. Mass fraction of indicated proteome subgroups on day 1 and day 9 of starvation (groups are based on GO cellular component annotation and quantified using iBAQ). Numbers inside sectors indicate mass fraction. Numbers outside sectors indicate the relative change in fraction size between days 1 and 9. **e**,**f**. Box plots of fold change in protein amount (normalized to total protein) between day 1 and day 9 for individual proteins in indicated groups (quantified using LFQ). Statistics: significance of comparison between groups (robust one-way ANOVA, adjusted post-hoc comparisons). ***p<0.001 **p<0.01. Only the most relevant comparisons are indicated. See Supplemental Table S1 for full statistics and precise p-values. **g**. Protein quantification by HiBit normalized to total protein and to values at t = 0. error bars: s.e.m. from 3 biological repeats. line: linear regression (*rpl-17:HiBiT* r^2^=0.94, *vit-2:HiBiT* r^2^=0.79). shaded area: 95% CI of linear fit. **h**. Mean volumetric protein density of GFP-fluorescence over starvation time. Shaded area: 95% CI. n = 146, 323 and 137 animals from 3 to 4 biological repeats for *lmn-1:gfp, daf-21:gfp* and *rpl-29:gfp*. **i**. Images of indicated strains after 0h and 55h starvation in agarose chambers. White: animal outline. Color scale: GFP intensity with equal contrast adjustment for the same genotypes. Animals were computationally straightened for display.

We confirmed these findings by LFQ-based fold change measurements on a per-protein basis, followed by regrouping: ribosomal proteins, on average, declined more strongly than other cytoplasmic proteins (Fig. 1e, Fig. S1a-b), as did translation-related proteins in general (Fig. 1f). Notably, mitochondrial ribosomal proteins also declined more than the average mitochondrial proteome (Fig. S1a). This intra-mitochondrial bias may explain the stronger decline of total mitochondrial proteins in iBAQ data compared to LFQ, as our iBAQ analysis weighed individual proteins by their mass abundance. Yolk proteins also declined rapidly, although this change was not statistically significant, given the small number of proteins (n=3) detected in this group (Fig. 1e).

Daily quantification of selected individual proteins through luminescence-based detection by HiBiT ^21^ (Fig. 1b) confirmed these results: endogenously HiBit-tagged ribosomal protein RPL-17 and yolk protein VIT-2, declined at 7.0% and 10.9% per day normalized to total protein (Fig. 1g).

To directly measure protein decline in individual animals, we performed live imaging using an endogenously tagged homozygous RPL-29:GFP insertion and integrated transgenes expressing non-ribosomal cytoplasmic DAF-21:GFP^22^ and nuclear lamin LMN-1:GFP^23^. In arrayed agarose-based chambers ^24^, we starved L1 larvae after hatching from eggs and imaged each individual hourly for 55 hours. Consistent with proteomics-based measurements, the fluorescence density (fluorescence per volume) of the ribosomal protein RPL-29:GFP declined during starvation, nuclear LMN-1:GFP increased, and cytosolic DAF-21:GFP remained approximately constant (Fig. 1h,i & Fig. S1d-e). Fluorescence decline was not due to photobleaching, which was negligible (Fig. S1c).

Taken together, different cytoplasmic proteins decline at different rates during starvation, resulting in a global remodeling of the organismal proteome and a reduced representation of growth-related proteins, including ribosomal and translation-associated proteins, as well as mitochondrial proteins and mitochondrial ribosomes.

### Ribosomal protein decline is autophagy-dependent

To test if ribosomal protein decline requires autophagy, we measured RPL-29:GFP in starving L1 larvae mutated for *atg-18*/WIPI or *unc-51*/ULK1 ^25,26^ over 55 hours by live imaging. All strains remained viable during the imaged period, as validated by the absence of death-induced swelling (Fig. S2d). Unlike for wild type animals, no decline in RPL-29:GFP fluorescence per volume was detectable in these mutant strains (Fig. 2a). A residual decline in total GFP fluorescence of 5% to 10% was detectable in both mutants when fluorescence was analyzed without volume normalization (Fig S2a,c). However, this decline in GFP was matched by a proportional decline in body volume (Fig. S2b), such that RPL-29:GFP concentration remained constant over time.

**Figure 2.**
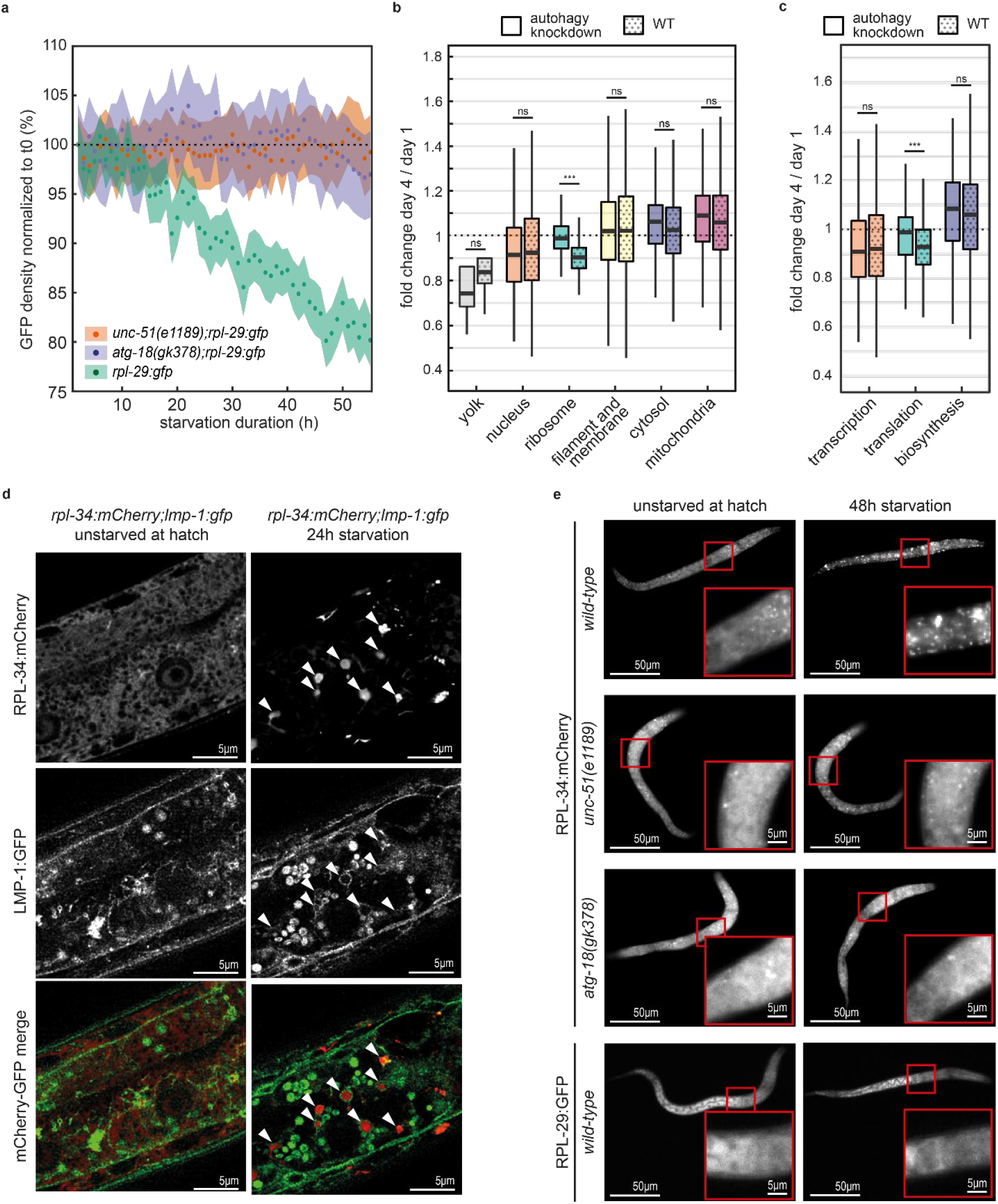
Ribosomal protein decline is autophagy-dependent. **a**. RPL-29:GFP fluorescence concentration (fluorescence per volume) in wild type and in autophagy deficient animals during starvation. Mean and 95% CI of the mean for n = 283, 299, 101 animals from 3 biological repeats for *unc-51(e1189)* and *atg-18(gk378)* and 2 biological repeats for wild type. **b**. Fold change from day 1 to day 4 of starvation (normalized to total protein) for ATG-18 depleted and control animals measured by TMT-proteomics. Box plots show fold change of individual proteins in indicated groups. n = 3 biological repeats. *** p < 0.001. See Supplemental Table 1 for precise p-values **c**. same as b., but for proteins associated with indicated GO terms. **d**. Co-localization of RPL-34:mCherry and lysosomal membrane associated protein LMP-1:GFP at hatching and after 24h of starvation. Arrowheads: ribosomal foci, surrounded by LMP-1:GFP. **e**. RPL-34:mCherry and RPL-29:GFP in wild type and autophagy-deficient individuals at hatch and after 48h starvation. Contrast settings are equal for the same genotype. Focus formation is autophagy and starvation dependent, and only occurs for acid-insensitive mCherry, but not for acid-sensitive GFP.

We further validafted the autophagy-dependence of ribosomal protein decline by tandem-mass-tag (TMT) labeled proteomics by comparing animals depleted for ATG-18 by auxin inducible degradation (AID) ^27^ to control animals (detecting 5880 proteins). While animals with intact autophagy showed the expected decline in ribosomal (-9%), and translation-related proteins (-7%) after 4 days, this decline was significantly diminished in ATG-18 depleted animals (Fig. 2b,c, p < 10^-4^). Differences between ATG-18 AID and control animals were not significant for other protein groups, likely because the decline in the more stable proteins over short starvation periods was not sufficiently pronounced to be reliably detectable by mass-spectrometry. Starvation for longer than 4 days was not feasible due to declining viability of the ATG-18 depleted animals (Fig. S2h).

To assess the subcellular fate of ribosomal proteins, we endogenously tagged RPL-34 and RPL-22 with mCherry, which, unlike GFP, remains fluorescent in acidic lysosomes ^28–30^. Upon starvation, RPL-34:mCherry and RPL-22:mCherry formed bright foci surrounded by lysosomal membrane marker LMP-1::GFP (Fig. 2d, Fig. S2e). These foci were less pronounced or entirely absent in *atg-18* and *unc-51* mutants (Fig. 2d, quantified in Fig. S2f). No foci were detected for a ribosomal protein tagged with an acid-sensitive fluorophore (RPL-29:GFP).

We conclude that the starvation-induced decline of ribosomal proteins is strongly autophagy-dependent, and that ribosomal proteins are delivered to lysosomes. Proteomics and live imaging show that any residual decline in ribosomal proteins scales proportionally with animal volume and proteome shrinkage, suggesting little to no disproportionate ribosome loss when autophagy is impaired.

### Ribosomal protein amount after starvation predicts recovery growth rate

To test whether ribosomal protein levels at the end of starvation relate to the recovery speed, we quantified RPL-29:GFP levels at the end of starvation and the growth rate during feeding for the same individuals by live imaging (Fig. 3a). Larval molts (M1 to M4) were detected by the halt in volume growth prior to cuticular ecdysis (Fig. 3b) at high temporal precision ^24,31^.

**Figure 3.**
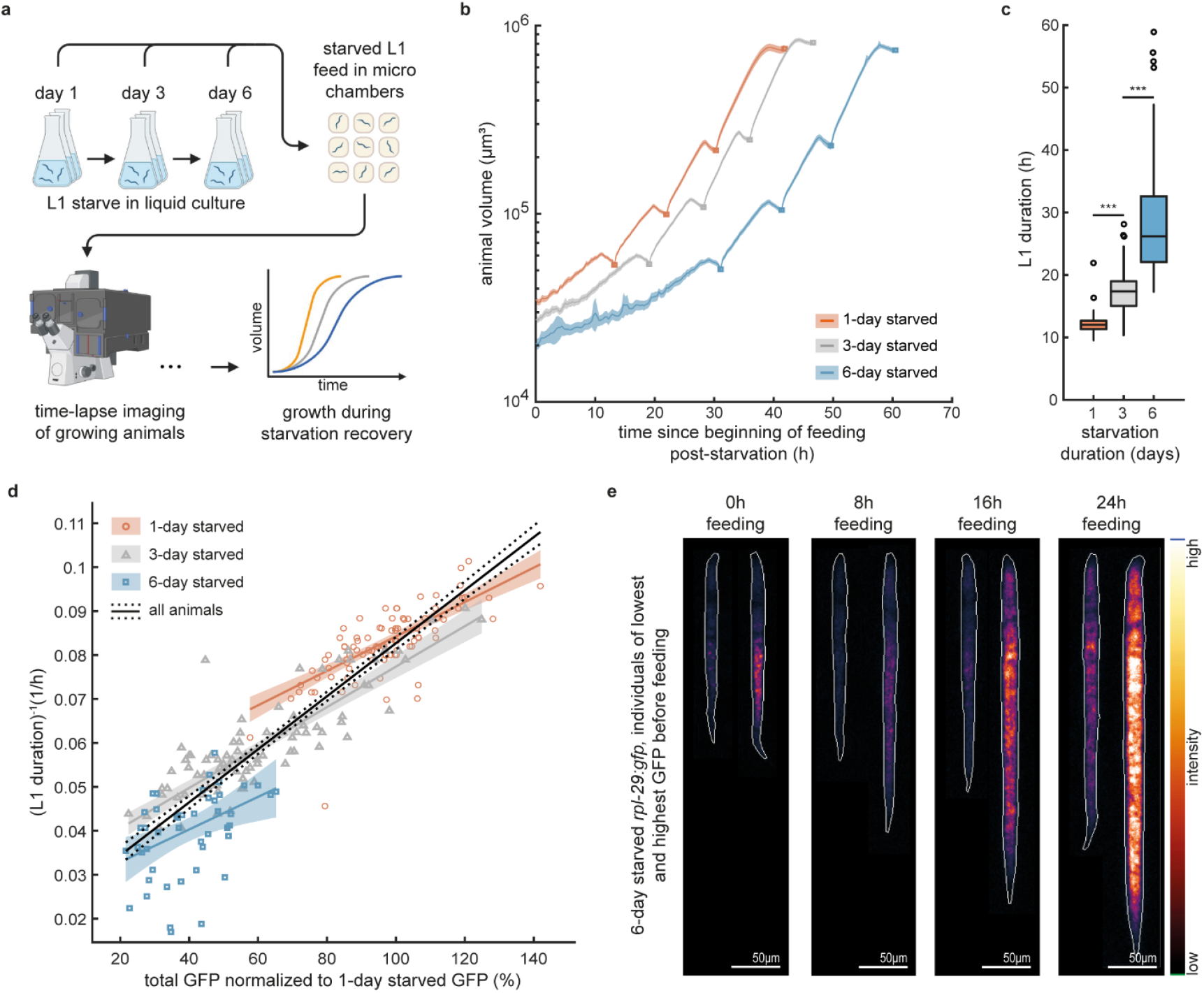
Ribosomal protein level predicts recovery speed. **a**. Experimental design of starvation recovery measurements. **b**. Mean animal volume during post starvation feeding after 1, 3 and 6-day starvation for n = 67, 66 and 43 indivduals. Squares: molts between larval stages. Shaded area: 95% CI of the mean. Means were computed by aligning individual trajectories to the molts prioir to averaging, followed by re-scaling to the mean larval stage duration. **c**. Box plots of L1 duration after 1, 3 and 6-day starvations for 89, 89 and 71 individuals, respectively. Significance between group means (two sample t-test): ***p<0.001. Precise p-values in Table S2 **d**. RPL-29:GFP fluorescence vs. the inverse of the L1 larval stage duration after recovery from 1, 3 and 6-day starvation for 85, 83 and 44 individuals. Lines: linear correlation with 95% CI. Black: all starvation durations pooled (r^2^=0.85). blue, gray, red: separated by starvation duration (r^2^=0.62, r^2^=0.76 and r^2^=0.19 for 1, 3, and 6 days). **e**. Images of the individuals expressing the highest and the lowest RPL-29:GFP after 6-day starvation at indicated times of feeding. Contrast is matched for animals at the same time of feeding. White: body outline. Color scale: GFP intensity. Animals were computationally straightened for display.

RPL-29:GFP progressively declined over 6 days of starvation (Fig. S3c), whereas the duration of the L1 stage during post-starvation refeeding progressively increased ^32,33^ (Fig. 3b, d). We observed substantial heterogeneity in the remaining RPL-29:GFP fluorescence among individuals, which was highly correlated with the recovery speed upon refeeding (r^2^=0.87, Fig. 3d, e). This correlation occurred across different durations of starvation, as well as across individuals exposed to the same duration of starvation (Fig. 3d). Thus, regardless of the starvation conditions experienced, the remaining RPL-29:GFP after starvation predicted the time a given individual needed to complete the first larval stage with high accuracy.

Starved animals recovered ribosomes to near normal levels during the first larval stage (Fig. S3c). Consistently, starvation duration had a little to no effect on larval stage durations, growth rates, ribosome expression, and body volumes from the second larval stage onwards (Fig. 3b, Fig. S3c-d). While we also detected a significant correlation between L1 duration and animal volume, this correlation was weaker than that with total RPL-29:GFP (r^2^ = 0.67 vs. r^2^ = 0.85, Fig. S3a) and a significant correlation (r^2^=0.64) also occurred when normalizing RPL-29:GFP by volume (Fig. S3b).

Together, these results establish ribosome content after starvation as a strong predictor for recovery speed, linking molecular resource levels to whole animal growth performance.

### Hyperactivation of the mTORC1 activator RAGA-1 reduces proteome remodeling and ribosomal decline

To test how metabolic signaling influences starvation-induced proteome remodeling and recovery speed, we analyzed a gain-of-function mutant of the mTORC1 activator *raga-1/*RagA that is stabilized in its active GTP-bound form^34^ (hereafter called RAGA-1^GTP^) ^35^ (Fig. 4a).

**Figure 4.**
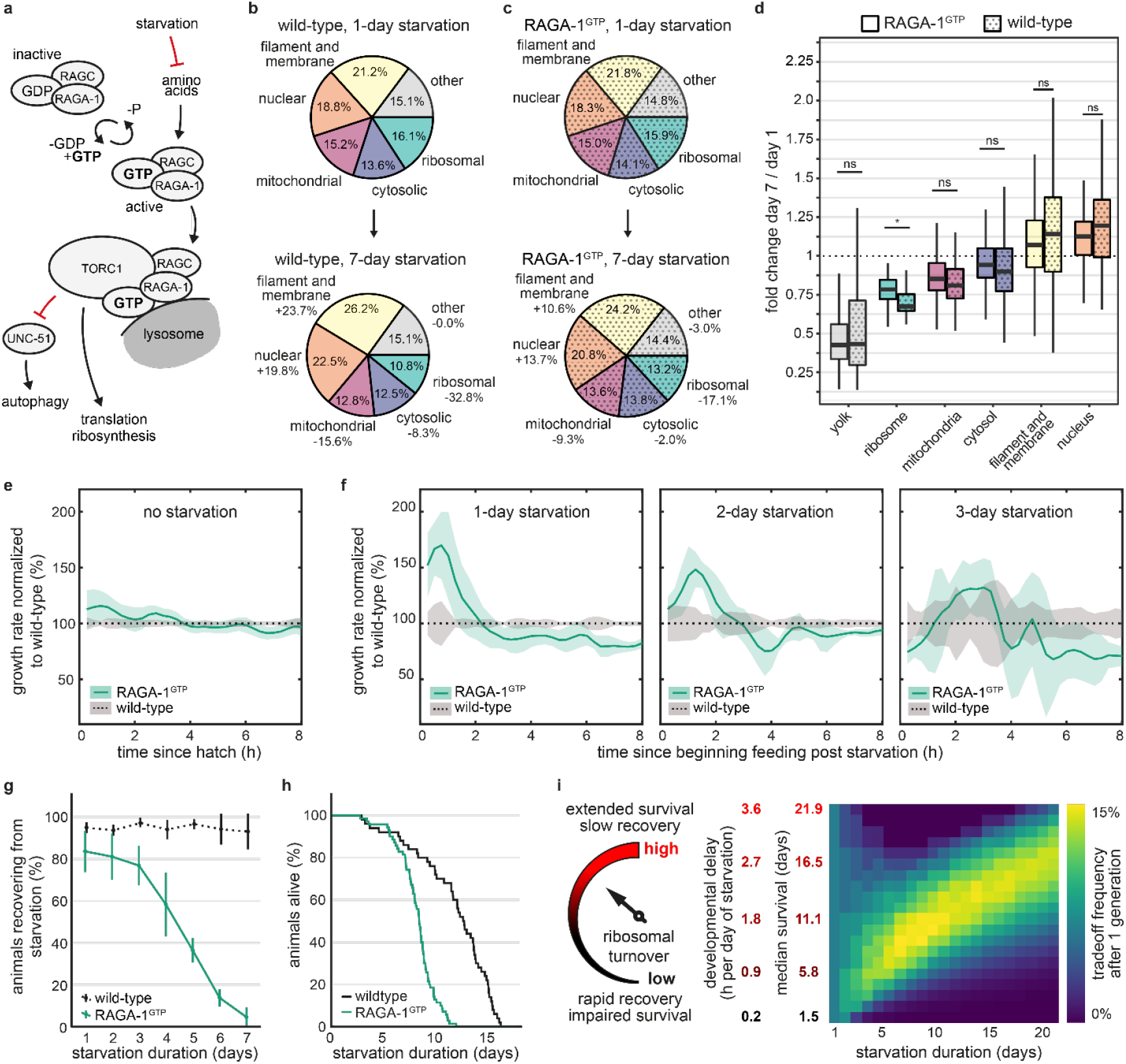
Reducing ribophagy shifts a tradeoff between starvation survival and recovery: **a**. Schematic of RAGA-1 in mTORC1 signalling. **b**,**c**. Quantification of proteome mass fractions on day 1 and 7 of starvation as described in Fig. 1, but for wild type and RAGA-1^GTP^ animals. Percentages inside sectors indicate group fraction size. Percentages outside sectors indicate proportional change in fraction size from the 1^st^ day to the 7^th^ day of starvation. **d**. Box plots of protein concentration fold change of individual proteins between day 1 and day 7 grouped by GO cellular component-terms for wild-type (solid fill) and RAGA-1^GTP^ (dotted fill). Statistics: differences between group means (robust two-way ANOVA, adjusted post-hoc comparison) * p = 0.011. **e**,**f**. Ratio of relative growth rate ((dV/dt)/V) between RAGA-1^GTP^ and wild type animals during *ad libitum* feeding after hatching (e) and after indicated duration of starvation (f) for the first 8 hours of feeding. Solid line: mean of 177 to 386 individuals across 2 to 5 biological repeats per strain and condition. Shaded area: 95% CI of the mean. **g**. Fraction of individuals that successfully recovered growth after indicated starvation duration. A total of 3467 wild type and 2621 RAGA-1^GTP^ individuals were sampled across 3 biological repeats. **h**. Starvation survival in agarose chambers. 48 wildtype and 70 RAGA-1^GTP^ animals. **i**. Proposed model: trade-off between starvation survival and starvation recovery is tuned by the rate of decline of ribosomes and possibly other proteins required for translation and growth. Heatmap shows simulated relative population growth for competing strains with different growth and survival balances (rows) under different durations of starvation (columns). The optimal compromise that maximizes selective fitness (yellow) depends on the duration of starvation.

We performed side-by-side quantitative label-free proteomics in wild-type and RAGA-1^GTP^ animals after 1, 4, and 7 days of starvation, a starvation duration during which viability of RAGA-1^GTP^ animals was not compromised compared to wild type (Fig. 4h, Fig. S5f). The mutation markedly reduced starvation-induced proteome changes with a stronger effect on ribosomal proteins (Fig. 4b-c, -32.8% ± 1.4% in wild type versus −17.1% ± 1.8% in RAGA-1^GTP^, Δ = -15.7%) compared to cytoplasmic (Δ = -6.3%) and mitochondrial proteins (Δ = -6.3%). Consequently, RAGA-1^GTP^ retained higher levels of ribosomal fraction than wild type (13.2% vs. 10.8%) by day 7 (Fig. 4b, c). LFQ-based analysis of individual proteins confirmed this pattern with ribosomal and translation-associated protein decline being significantly supressed in RAGA-1^GTP^ animals (Fig. 4d, S4a-c).

We validated the proteomics results by live imaging of RPL-29::GFP in wild-type and RAGA-1^GTP^ larvae. At hatching, RPL-29::GFP levels were ∼10% higher in RAGA-1^GTP^ than in wild type (Fig. S4d–e), consistent with mTORC1 hyperactivation. Over 55 hours of starvation, total fluorescence, fluorescence normalized to starting levels, and volumetric fluorescence density all remained higher in RAGA-1^GTP^ (Fig. S4d–g). Body volume declined at similar rates in both genotypes (Fig. S4h–i), indicating no major difference in overall biomass turnover.

Photobleaching assays showed negligible RPL-29::GFP production in both genotypes during starvation (Fig. S5a–d), excluding altered ribosome synthesis as the cause of slower decline in RAGA-1^GTP^. Fluorescence differences were also not attributable to expression of the GFP tag linked to the RAGA-1^GTP^ allele, which was near-negligible relative to RPL-29::GFP (Fig. S5e).

Together, these data show that the RAGA-1^GTP^ mutation dampens starvation-induced proteome changes, including the decline in ribosomal and other translation-associated components.

### RAGA-1^GTP^ impairs starvation survival and speeds up growth during recovery

In agreement with previous findings in mice ^36^, RAGA-1^GTP^ mutants showed reduced starvation resistance with more than 50% of the animals failing to recover growth after 5 days of starvation (Fig. 4g), and a sharp drop of viability after 9 days (Fig. 4h). To test if this reduced survival of extended starvation was paralleled by a faster recovery after short starvation, we measured the growth rate of wild-type and RAGA-1^GTP^ animals exposed to food after 1, 2, and 3 days of starvation by live imaging. Without starvation, growth rates were indistinguishable between genotypes (Fig. 4e). After 1 and 2 days of starvation, however, RAGA-1^GTP^ mutants grew significantly faster than wild-type during the first two hours of recovery (Fig. 4f, Fig. S6a-b,i-j). This advantage was transient and disappeared later in development when wild-type animals had restored ribosome levels to normal levels (Fig. S6c-f).

No growth benefit was observed after 3 days of starvation (Fig. 4f, Fig. S6b,i,j), possibly due to increased frailty of RAGA-1^GTP^ mutants after prolonged food starvation (Fig. 4g,h) or a smaller ribosome difference at this timepoint (Fig. S6e,f).

Together, these data show that RAGA-1^GTP^ accelerates growth recovery after short periods of starvation yet impairs growth and survival after longer starvation. A parsimonious explanation for this relation is that energy scavenged from the faster degradation of growth-related proteins in the wild-type supports long-term organismal integrity, but the need to re-synthesize these proteins incurs a cost during growth recovery ^37–42^ (Fig. 4i).

### Best compromise between recovery speed and survival depends on starvation duration

To illustrate when rapid recovery may be more advantageous than high starvation survival, we modelled population growth using realistic life-history parameters for *C. elegans*. In the model, starvation imposed a developmental delay that increased linearly with time, as observed experimentally ^43^. Starvation survival followed a Hill function with half-point matching measured data (Fig. 4h), and the survival-recovery tradeoff was represented by proportional coupling of these parameters (see methods).

We simulated competition between strains with different tradeoff strategies until the population expanded 20-fold. This simulation showed that a longer median survival at the cost of a longer delay developmental is beneficial for long durations of starvation, whereas a short delay and short survival are advantageous for brief starvation duration.

These results indicate that the temporal pattern of food availability in an environment influences the optimal balance between growth and survival. Altering the rate of autophagy-dependent proteome remodeling may adjust this balance.

## Discussion

In environments where nutrient availability fluctuates, animals must withstand extended periods without food and resume growth effectively when feeding resumes. We identify starvation-induced proteome remodeling as a molecular mechanism tightly connected to this balance.

Growth-related proteins, including ribosomal components, decreased in abundance more rapidly than other major protein groups during starvation (Fig. 1). Consistent with ribosomes acting as a limiting factor for protein synthesis and growth during recovery from starvation, animals with lower residual ribosomal protein levels after starvation required more time to complete the L1 stage (Fig. 3). Our results thereby connect proteome-scale changes with organism-scale life-history traits by linking ribosomal protein abundance to recovery growth dynamics.

Manipulating metabolic signalling supported the importance of proteome remodelling in starvation survival and recovery (Fig. 4). A constitutively active form of RAGA-1^44^, an activator of mTORC1 signalling^35^, reduced starvation-induced proteome remodelling and ribosomal protein loss, accelerated early growth recovery from short starvation, and decreased survival after prolonged starvation. These trade-offs are consistent with a shift in the balance between preserving growth capacity and allocating resources for long-term survival.

Autophagy-deficient mutants dampened the absolute ribosomal protein decline and abolished the decline relative to other proteins, suggesting autophagy as a main driver of proteome remodelling during starvation (Fig. 2). While receptor-mediated autophagy can target specific cellular components^9–12,16^, it is unclear whether the ribosomal protein decline we observe here is directed by such a pathway, or arises from receptor-independent, autophagy-dependent processes ^45,46^. Proteasomal decay and residual protein synthesis during starvation may also contribute, but the strong suppression of proteome remodelling upon ATG-18 depletion (Fig. 2a-c) argues against these processes as primary mechanisms.

Although our live imaging experiments focused on ribosomal proteins, our proteomic data show that also other growth-related factors and non-ribosomal proteins decline disproportionately to the rest of the proteome. This coordinated turnover likely contributes to the overall recovery phenotype. Nevertheless, the strong near-linear relation between RPL-29:GFP and developmental rate, explaining 85% of the variance, suggests that ribosomal protein levels are a strong predictor for the growth rate. This strong correlation is reminiscent of “growth laws” described in microbes, in which protein allocation linearly and predictably scales with growth and survival rates ^47–49^. The coherence of our data with such growth laws suggests that core principles of proteome allocation may extend from microbes to multicellular systems.

In summary, we identify autophagy-dependent ribosomal protein decline as a major contributor to starvation-induced proteome remodelling and show that the proteome composition at the time of refeeding is strongly related to recovery dynamics in *C. elegans*. By quantitatively connecting molecular traits to organismal life-history traits, our work establishes a new paradigm for understanding the mechanisms and evolutionary implications of proteome resource allocation during nutrient stress.

## Materials and Methods

### C. elegans strains and maintenance

Animals were grown and maintained at 25°C using OP50-1 *E. coli* on nematode growth medium (NGM) according to standard procedures. The following strains were used in this study:

WBT100: *rpl-17(xe232)[rpl-17:HiBit]; xeSi296[peft-3::luc-gfp::unc-54 3’UTR, unc-119(+)] II* (this study)

WBT203: *vit-2(syb3768)[vit-2::GSSG::3xflag::GSSG::HiBiT] X* (this study)

GW8: *ygIs[lmn-1::GFP::lmn-1 unc-119(+)] X; unc-119(ed3) III* (^23^)

RBW2661: *hutSi2661 [hsp-90p::eGFPT::unc-54 3’UTR + Cbr-unc-119 (+)] II* (^22^)

PHX1945: *rpl-29(syb1945) [rpl-29:GFP]* (this study)

N2: Bristol N2 wild isolate

WBT257: *rpl-29(syb1945) [rpl-29:GFP] IV; unc-51(e1189) V* (this study, allele e1189 from CGC registered strain CB1189 by J. A. Hodgin lab, Oxford University)

WBT277: *rpl-29(syb1945) [rpl-29:GFP] IV; atg-18(gk378) V* (this study, allele gk378 from ^50^) PHX1880: *rpl-34(syb1880) [rpl-34:RPL-34:mCherry] IV* (this study)

PHX1903: *rpl-22(syb1903) [rpl-22:RPL-22:mCherry] II* (this study)

WBT230: *rpl-34(syb1880) [rpl-34:RPL-34:mCherry] IV; unc-51(e1189) V* (this study, allele e1189 from CGC registered strain CB1189 by J. A. Hodgin lab, Oxford University)

WBT234: *rpl-34(syb1880) [rpl-34:RPL-34:mCherry] IV; atg-18(gk378) V* (this study, allele gk378 from ^50^)

WBT343: *unc-119(ed3)III; pwls50[lmp-1p::lmp-1::GFP+Cb-unc-119(+)]; rpl-34(syb1880) [rpl-34:RPL-34:mCherry] IV* (this study, allele pwls50 from CGC registered strain RT258 by B. D. Grant lab, Rutgers University)

WBT64: *xeSi296 [Peft-3::luc::gfp::unc-54 3’UTR, unc-119(+)] raga-1(xe190) II* (this study, allele xeSi296 from ^51^, allele xe190 from ^34^)

HW1939: *xeSi296 [peft-3::luc::gfp::unc-54 3’UTR, unc-119(+)] II* (^51^)

WBT330: *raga-1(wbm40) [raga-1::AID::EmGFP] II; xeSi376ubs38[Peft-3::TIR1(*ubs38 *[F79G])::mRuby::unc-54 3’UTR, cb-unc-119(+)] III; rpl-29(syb1945) [rpl-29:GFP] IV* (this study, allele xeSi376ubs38 from ^52^)

WBT289: *raga-1(wbm40abt3) [raga-1(GF)::AID::EmGFP] II;* xeSi376ubs38*[Peft-3::TIR1(*ubs38 *[F79G])::mRuby::unc-54 3’UTR, cb-unc-119(+)] III; rpl-29(syb1945) [rpl-29:GFP] IV* (this study, allele xeSi376ubs38 from ^52^)

WBT231: *raga-1(wbm40abt3) [raga-1(GF)::AID::EmGFP] II* (this study)

WBT282: *hibit::aid::atg-18 V*.

WBT283: *hibit::aid::atg-18 V*.; *ieSi57 [eft-3p::tir1::mRuby::unc-54 3’UTR, cb-unc-119(+)] II*.

### Egg collection and starvation cultures

Animals were grown on NGM plates at 25°C for at least 3 generations prior to starvation. For mass spectrometry experiments, the peptone concentration in NGM was increased to 20mg/ml to reach sufficient numbers of individuals. Mixed populations were synchronized by bleaching and the eggs were allowed to develop into gravid adults on new plates. The adults were filtered with a cell strainer with 40µm mesh size (Pluriselect) to remove younger animals and incubated in 35mM serotonin in S-basal in a rotating cradle at 25°C for 1.5h. Eggs were harvested and separated from adults with a 40µm mesh, and the eggs were filtered a 20µm mesh (Neolab) to remove hatched L1 larvae. The purified eggs were then either loaded into agarose chambers for timelapse imaging or allowed to hatch overnight in S-basal (without cholesterol and ethanol) in a rotating cradle at 25°C at a concentration of 10 eggs/µl. The following day, the hatched L1 larvae were filtered with 20µm mesh to remove unhatched eggs. The L1 larvae were then starved in S-basal (without cholesterol and ethanol) in a rotating cradle at 25°C at a density of 10 L1/µl for the remainder of the starvation duration.

For mass spectrometry experiments, the synchronized gravid animals were bleach synchronized instead of serotonin treated to obtain eggs. The eggs were allowed to hatch overnight in S-basal (without cholesterol and ethanol) in an orbital shaker at 25°C at a concentration 10 eggs/µl. The hatched L1 larvae were filtered with a 20µm mesh to remove unhatched eggs. The L1 larvae were starved in sterile S-basal cultures (without cholesterol and ethanol) in orbital shaker at 25°C at 10 L1/µl for the remainder of the starvation duration.

### Label-free mass spectrometry and analysis

Starving L1 larvae were collected (10^6^ animals/sample) from triplicate cultures (S-basal without cholesterol and ethanol at 25°C in orbital shaker at 10 animals/µl) and resuspended into 700µl PBS without additives. The pellets were flash frozen by pipetting dropwise into liquid nitrogen. Frozen beads were powdered using cryogenic SPEX SamplePrep 6775 Freezer/Mill and stored as frozen powder at -80°C.

Powdered pellets were dissolved in 1% sodium deoxycholate buffer containing 50 mM Tris-HCl pH 8. Samples were treated with 1 mM DTT and alkylated using 5.5 mM iodoacetamide for 10 min at RT. Samples were in-solution digested with trypsin (Promega). Tryptic peptides were purified by STAGE tips and LC-MS/MS measurements were performed on a QExactive plus and an Orbitrap Exploris 480 mass spectrometer coupled to EasyLC 1200 nanoflow-HPLCs (all Thermo Scientific) using data-dependent acquisition. MaxQuant software (version 1.6.2.10 & 2.0.1.0) ^53^ was used for analyzing the MS raw files performing peak detection, peptide quantification and identification using a Uniprot *C. elegans* database, November 2017. Carbamidomethylcysteine was set as fixed modification and oxidation of methionine was set as variable modification. The MS/MS tolerance was set to 20 ppm and four missed cleavages were allowed for Trypsin/P as enzyme specificity. Based on a forward-reverse database, protein and peptide FDRs were set to 0.01, minimum peptide length was set to seven, and at least one unique peptide had to be identified. MaxQuant results were analysed using Perseus software (version 1.6.2.3)^54^. iBAQ and LFQ values were used for expression proteomics analyses. iBAQ values of the respective samples were median normalized. Only proteins with valid values in all three biological replicates in respective groups were considered for further analysis.

iBAQ and LFQ data were analyzed using R-studio (v4.3.2). Mass fractions were determined by multiplying iBAQ quantification of each detected protein by its molecular weight. Resulting mass values were grouped and summed based on GO terms and divided by the total iBAQ mass. Molecular weights and GO terms were acquired from UniProtKB by list search of protein IDs reported by MaxQuant. Fold changes of individual proteins were grouped based on whether the GO terms associated with the protein matched category presented in Fig. 1d-f and 4d (e.g. “ribosome”). For figures shown in main text, each protein was allowed to appear in only one group (see Table S1 for list of proteins in each group).

### Tandem-mass-tag (TMT) Proteomics and analysis

Sample collection: Animals with an aid tag endogenously inserted at the *atg-18* locus and expressing Tir1 under the ubiquitously active *eft-3* promoter (strain WBT283, for ATG-18 depletion) and without Tir1 expression (WBT282, control) were grown to adulthood in the absence of auxin. Adult animals were bleached as described above to prepare eggs. Both strains were allowed to hatch overnight without auxin, and auxin (dissolved in DMSO at a concentration of 250mM) was added to a final concentration of 0.5mM. On days 0, 2, and 4, a sample of 100’000 animals was collected from each culture for mass spectrometry and a small subsample was inspected for viability (based on animal motility). Animals were lysed in 100μl lysis buffer (PBS with 0.25% Triton-X100 and proteinase inhibitors (Sigma)) by three consecutive freeze-thaw cycles and subsequent sonication in a bioruptor (10 min, 30s on/off cycles, 4°C). Soluble fraction was retrieved after centrifugation 16’000g for 10 minutes at 4°C. Protein concentration was determined by microBCA and adjusted to 20μg in 60μl per each sample. Three separate biological repeats of L1 larvae, stemming from separate parental cultures, were collected for each condition and day.

Protein samples were subjected to the SP3 protocol ^55^ conducted on the KingFisher Apex™ platform (Thermo Fisher). For digestion, trypsin was used in a 1:20 ratio (protease:protein) in 50 mM N-2-hydroxyethylpiperazine-N-2-ethanesulfonic acid (HEPES) supplemented with 5 mM Tris(2-carboxyethyl)phosphine hydrochloride (TCEP) and 20 mM 2-chloroacetamide (CAA). Digestion was carried out for 5 hours at 37°C.

Up to 10 µg of peptides were labeled using TMTpro™ 18plex reagent as previously described ^56^. Briefly, 0.5 mg of TMT reagent was dissolved in 45 µL of 100% acetonitrile. Subsequently, 4 µL of this solution was added to each peptide sample, followed by incubation at room temperature for 1 hour. The labeling reaction was quenched by adding 4 µL of a 5% aqueous hydroxylamine solution and incubating for an additional 15 minutes at room temperature. Labeled samples were then combined for multiplexing, desalted using an Oasis® HLB µElution Plate (Waters) according to the manufacturer’s instructions, and dried by vacuum centrifugation.

Offline high-pH reversed-phase fractionation ^57^ was carried out using an Agilent 1200 Infinity high-performance liquid chromatography (HPLC) system, equipped with a Gemini C18 analytical column (3 μm particle size, 110 Å pore size, dimensions 100 x 1.0 mm, Phenomenex) and a Gemini C18 SecurityGuard pre-column cartridge (4 x 2.0 mm, Phenomenex). The mobile phases consisted of 20 mM ammonium formate adjusted to pH 10.0 (Buffer A) and 100% acetonitrile (Buffer B). The peptides were separated at a flow rate of 0.1 mL/min using the following linear gradient: 100% Buffer A for 2 minutes, ramping to 35% Buffer B over 59 minutes, increasing rapidly to 85% Buffer B within 1 minute, and holding at 85% Buffer B for an additional 15 minutes. Subsequently, the column was returned to 100% Buffer A and re-equilibrated for 13 minutes. During the LC separation, 48 fractions were collected. These were pooled into twelve fractions by combining every twelfth fraction. The pooled fractions were then dried using vacuum centrifugation.

The samples were then measured on an UltiMate 3000 RSLCnano LC system (Thermo Fisher Scientific) equipped with a trapping cartridge (µ-Precolumn C18 PepMap™ 100, 300 µm i.d. × 5 mm, 5 µm particle size, 100 Å pore size; Thermo Fisher Scientific) and an analytical column (nanoEase™ M/Z HSS T3, 75 µm i.d. × 250 mm, 1.8 µm particle size, 100 Å pore size; Waters) coupled to an Orbitrap Fusion™ Lumos™ Tribrid™ mass spectrometer (Thermo Fisher Scientific) . Samples were trapped at a constant flow rate of 30 µL/min using 0.05% trifluoroacetic acid (TFA) in water for 6 minutes. After switching in-line with the analytical column, which was pre-equilibrated with solvent A (3% dimethyl sulfoxide [DMSO], 0.1% formic acid in water), the peptides were eluted at a constant flow rate of 0.3 µL/min using a gradient of increasing solvent B concentration (3% DMSO, 0.1% formic acid in acetonitrile). Peptides were introduced into an Orbitrap Fusion™ Lumos™ Tribrid™ mass spectrometer (Thermo Fisher Scientific) via a Pico-Tip emitter (360 µm OD × 20 µm ID; 10 µm tip, CoAnn Technologies) using an applied spray voltage of 2.2 kV. The capillary temperature was maintained at 275 °C. Full MS scans were acquired in profile mode over an m/z range of 375–1,500, with a resolution of 120,000 at m/z 200 in the Orbitrap. The maximum injection time was set to 50 ms, and the AGC target limit was set to ‘standard’. The instrument was operated in data-dependent acquisition (DDA) mode, with MS/MS scans acquired in the Orbitrap at a resolution of 30,000. The maximum injection time was set to 94 ms, with an AGC target of 200%. Fragmentation was performed using higher-energy collisional dissociation (HCD) with a normalized collision energy of 34%, and MS2 spectra were acquired in profile mode. The quadrupole isolation window was set to 0.7 m/z, and dynamic exclusion was enabled with a duration of 60 seconds. Only precursor ions with charge states 2–7 were selected for fragmentation. Raw files were converted to mzML format using MSConvert from ProteoWizard, using peak picking, 64-bit encoding and zlib compression, and filtering for the 1000 most intense peaks. Files were then searched using MSFragger in FragPipe (22.1-build02) against FASTA database UP000001940_CaenorhabditisElegans_BristolN2_ID6239_19832entries_05042024_dl17062024 containing common contaminants and reversed sequences. The following modifications were included into the search parameters: Carbamidomethylation (C, 57.0215), TMTpro (K, 304.2072) as fixed modifications; Oxidation (M, 15.9949), Acetylation (protein N-terminus, 42.0106), TMTpro (peptide N-terminus, 304.2072) as variable modifications. For the full scan (MS1) a mass error tolerance of 20 PPM and for MS/MS (MS2) spectra of 20 PPM was set. For protein digestion, ‘trypsin’ was used as protease with an allowance of maximum 2 missed cleavages requiring a minimum peptide length of 7 amino acids. The false discovery rate on peptide and protein level was set to 0.01.

For the data analysis the raw output files of FragPipe (protein.tsv files files) were processed using the R programming environment (ISBN 3-900051-07-0). Initial data processing included filtering out contaminants and reverse proteins. Only proteins quantified with at least 2 razor peptides (with Razor.Peptides >= 2) were considered for further analysis. 5879 proteins passed the quality control filters. In order to correct for technical variability, batch effects were removed using the ‘removeBatchEffect’ function of the limma package ^58^ on the log2 transformed raw TMT reporter ion intensities (‘channel’ columns). Subsequently, normalization was performed using the ‘normalizeVSN’ function of the limma package (VSN - variance stabilization normalization - ^59^). Missing values were imputed with the ‘knn’ method using the ‘impute’ function from the Msnbase package ^60^. This method estimates missing data points based on similarity to neighboring data points, ensuring that incomplete data did not distort the analysis. Analysis and grouping of fold changes was equivalent to that used for LFQ analysis.

### HiBiT measurements

Starving L1 larvae were collected (3000 animals/sample) from triplicate cultures into 100µl lysis buffer (PBS pH8, 0.25% triton, protease inhibitors (Sigma P8340)), and freeze-thawed three times in liquid nitrogen and 37°C water bath. Samples were lysed by sonication (Bioruptor) at 4°C and diluted with 600 µl PBS prior to HiBiT assay.

After lysis all pipetting was conducted using motorized pipettes (Sartorius) and a pipetting robot (Opentron) to minimize experimental variation. HiBiT intensity was measured daily in three biological replicates (3 measurements x 3 wells x 3 replicates / day) in 384-well format using a HiBiT quantification kit (Promega Nano-Glo). 5µl HiBiT reaction mixture (1:100 LgBiT protein and 1:50 HiBiT lytic substrate in HiBiT lytic buffer) was combined with 5µl sample. Plates were shaken at room temperature for 5min prior to measuring luminescence on a luminometer (SpectromaxL).

Protein concentration in each sample was measured using micro-BCA kit (Pierce) in three biological replicates (3 measurements x 3 wells x 3 replicates / day) in 96-well format. 140ul sample was combined with 140ul BCA reagent (25:24:1 mix of reagents MA:MB:MC) at 4°C. The plates were shaken for 5min, incubated stationary in 37°C incubator for 2h, and measured with Spectromax340 562nm in endpoint mode.

Data analysis was done in R (v4.3.2). Background signals were subtracted (BCA reagent with lysis buffer for micro-BCA, N2 worm lysate with reaction mixture for HiBiT) and HiBiT measurements were normalized with total protein BCA measurements to acquire HiBiT fraction. The fractions were normalized again by the average of the 3 biological replicates at t=0 and a linear model was fitted to the normalized values.

### Live imaging in agarose-based chambers

Arrayed agarose chambers were manufactured by stamping a microscopic chamber pattern into gel of 4.5% agarose in S-basal using a PDMS template (Wunderlichips) based on Turek et al. ^61^, incorporating the modifications by Stojanovski et al. ^24^. Animals were retained isolated in separate chambers and imaged every hour for starvation experiments, and every 15 minutes for post-starvation growth experiments. Chamber dimensions were 270x270x20µm for L1 starvation experiments in Fig. 1, Fig. 1S, Fig. 2 and Fig. 2S. Chamber depth was decreased to 10µm for L1 starvation experiments in Fig. 4S and for all L1 starvation recovery experiments in Fig. 4 to improve image focus, accuracy of segmentation, and volume estimates. Chamber dimensions were increased to 600x600x20µm for post-starvation growth experiments in Fig. 3 and Fig. S3 to accommodate animal growth at L3 and L4 stages.

For starvation experiments, the chambers were manufactured without adding cholesterol and ethanol to the agarose. 0.003% Triton was added to the eggs to reduce clumping and the eggs were pelleted. The egg pellet was gently broken up before pipetting 5µl concentrated eggs onto freshly manufactured chambers. 2-fold or older eggs inside a droplet of S-basal without cholesterol or ethanol were then individually guided to chambers using an eyelash pick.

For post-starvation growth experiments, chambers were filled with OP50-1 *E. coli* by scraping an *E. coli* lawn off a standard NGM plate and spreading it onto the chambers with a piece of 3% agarose gel prior to loading animals as described by Stojanovski et al. ^24^. In Fig. 4, the agarose of the chambers contained 0.05% DMSO. Starved L1 larvae were pipetted next to the array of chambers and then moved individually to chambers using an eyelash pick. Chamber array was prevented from drying during loading by pipetting 20µl deionized water around the outer perimeter of the chamber array.

After loading animals into the chambers, the agarose chambers were inverted onto dishes with gas permeable polymer bottom of high optical quality (Ibidi) and surrounded by 3% low melting temperature agarose. The assembly was covered with thin layer of PDMS to prevent evaporation and the dish was closed and sealed with parafilm and used for imaging.

A Nikon Ti2 wide-field epifluorescence microscope equipped with a SpectraX light source (Lumencor) and a Prime BSI camera (Photometrics) was used for all live imaging experiments. A 10x 0.45NA objective was used in Fig. 3b-e, Fig. S3a-d, Fig. S4e-g, and Fig. S6. A 20x 0.75NA objective was used in Fig. 1h-I, Fig. S1c-e, Fig. 2a, Fig. S2a-f, and Fig. S5a-h. Effective pixel size in the sample plane was 0.65 and 0.325µm, respectively. Software auto-focus of NIS Elements Software (Nikon) was used to find the central focal plane in the GFP or mCherry channel of the ribosomal markers using single band pass filters. For all experiments, the temperature was maintained at 25 °C using an incubator encapsulating the entire microscope (life imaging services).

### Confocal imaging of RPL-22:mCherry foci

In Fig. 2d, a Nikon CREST X-light spinning disk confocal microscope was used to image L1 animals immobilized on agarose pads by 10mM Levamisole and a coverslip. Animals were imaged at hatch or after 24h starvation. A 100x 1.45NA oil immersion objective was used to capture Z-stacks around manually set focus plane with a single band pass filters for mCherry and GFP using 200ms exposure. Images were deconvoluted using Huygens software.

### Detection of animals and animal fluorescence in agar chambers

A Matlab script was used to process raw images as described ^24^. Sobel algorithm using the edge function of Matlab (R2022b) was used to segment the animals from raw fluorescent images. Detected outline endpoints closest to each other were connected to close gaps in the animal outline. From the outline a binary mask was created to separate animals from background. For L1 starvation experiments in Fig. 1 and Fig. S1 a U-net ^62^ network was trained to segment the binary mask of animals from raw brightfield images. Segmentation was performed in brightfield channel to allow for the segmentation of non-fluorescent N2 control animals. The Sobel-based masks were used as the U-net training data. For both methods animal fluorescence was measured from the area covered by the mask in the fluorescent channel. For Fig. 1h-I, N2 animals were imaged in parallel and their auto fluorescent signal was subtracted from all measurements due to the comparatively weak fluorescence signal of LMN-1:GFP. For Figs. 2a and 3d, only the background signal from agar or bacteria was subtracted, as the tissue autofluorescence was negligible compared to the fluorescence of RPL-29:GFP (Fig. S5e).

The masks were computationally straightened using a custom made Matlab function ^24^. Straightened masks were classified as either ‘egg’, ‘worm’ or ‘error’ using a decision tree-based classifier that was trained on a small subset of manually annotated masks. The classifier was an ensemble of 20 bootstrap aggregation decision trees (TreeBagger function of Matlab) that was trained on features length, standard deviation of width, cv of width, maximal width, median width, maximal width/median width, volume, volume/length, and entropy of width ^24^. All experiments were classified with the same classifier, except for experiments with autophagy deficient animals, whose occasionally irregular outline necessitated training of a separate classifier for reliable error detection. Values derived from images with errors or missing animals were excluded from further analyses.

### Computation of volume and growth rate

The volume of the animals in agarose chambers was derived from the straightened binary image masks assuming rotational symmetry ^63^. For growing animals, larval stage transitions were determined based on the timing of the maximum of the second time derivative of the logarithm of animal volume. Using a graphical user interface created in Matlab ^24^, the ecdysis timings were corrected manually based on plateaus in animal growth curves caused prior to ecdysis. To minimize measurement error, for Fig. 3d and Fig. S3a,b, the fluorescence and volume at t0 was estimated by fitting a linear model to the first 20 timepoints and the predicted value at t0 was plotted. Individuals for which less than 6 data points were available in the first 20 timepoints were excluded. For Fig. 3c, the volume measurement was scaled to the larval development by linearly interpolating the volume at 100 points per larval stage. The interpolated values are plotted over the mean larval stage duration, such that the beginning and end of each larval stage are aligned on the x-axis. The same scaling was used for fluorescence in Fig. S3c.

The rate of growth during starvation recovery for Fig. 4e-f was calculated in Matlab by smoothing every individual animal’s volume with the fit function’s smoothing spline option using a smoothing parameter of 0.0001. Fits were manually inspected to ensure that the resulting volume interpolations are appropriate, after which volume was interpolated at a time resolution of 1 minute. Volume growth rate by minute was calculated as the local slope of the smoothed volume curve by using gradient function. Individuals’ growth rates were normalized to the mean growth rate of the control condition, and a mean of normalized growth rates was determined for every replicate chamber array.

### Photobleaching based detection of GFP synthesis

L1 larvae starved for 16h in agarose chambers were imaged hourly for three hours before exposure to a 477nm laser (Celesta light engine, Lumencor) for 8 seconds at 100% laser power until approximately 60% of GFP signal had been bleached. Viability was not affected based on motility. Animals were subsequently imaged for 20 hours, the auto-fluorescence was subtracted using N2 animals as control, and the fluorescence signal was normalized by the first timepoint after photobleaching (Fig. S5b-c).

Data analysis: The net change in total fluorescence of GFP (*g*) is the difference in its production with rate *p* and degradation with rate constant *d*, such that:

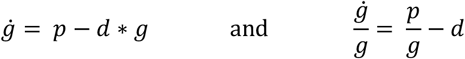

Thus, for p > 0, the relative rate of change in GFP 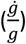 would increase after photobleaching, assuming that the rate constant d is unaffected by photobleaching. Conversely, if 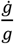 is unaffected by photobleaching (as observed in Fig. S5b,d), *p* must be negligible.

### Starvation recovery and survival assays

The ability of animals to recover growth after starvation (Fig. 4g) was measured daily at 25 °C by placing a total of 50-150 starving L1 larvae onto the bacterial lawn of fresh NGM plates from 3 independent cultures and counted immediately. After 2 or 3 days of growth (for shorter and longer than 4 day-starvation, respectively) animals that had exited L1 (based on their larger size) were considered to have resumed development and recovered from starvation.

Starvation survival (Fig. 4h) was determined by placing a minimum of 36 eggs per strain into agarose chambers without food and imaging every 17 min for 3 weeks at 25 °C. Worms were computationally detected as described above. Time of death was determined as the time of permanent loss of motility that was followed by death associated swelling (Fig. S2d).

### Population Growth Model

Population growth for different tradeoff strategies was computed using numerical simulations in Matlab(R2022a) by calculating the relative fraction of animals among different strategies after different times of starvation during one generation. Survival *s* after time *t* of starvation was described by a Hill function with Hill coefficient n=4 and half-point *σ* .

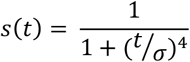

The lag time τ was determined by a linear relation τ = δ * t, where δ is the additional lag time per time of starvation. The tradeoff was model by the linear coupling *σ* = *k* ∗ *δ*. Parameter k was set to 56 days. 20 strains with *δ* being linearly spaced between 0.01 and 0.15 were computationally competed against each other, starting with a uniformly distributed population. This parameter range corresponds to a median survival *σ* spanning between 1.5 days and 22 days, and a lag times τ after 7 days of starvation between 1.7 hours and 25 hours, centered around the experimentally observed values shown in Fig. 3. Reproduction was kept equal among all strains, starting after 40 hours plus the respective lag time τ at a constant reproduction rate of 6 progeny per hour for 48 hours. The population was allowed to grow until it multiplied 20 times to simulate food exhaustion. The data shown in Fig. 4i is the relative frequency of each strain in the entire population at the end of the first starvation/recovery cycle.

### Statistical analysis of proteomic data

Proteins were assigned to functional groups based on Gene Ontology cellular component and biological process terms. To maintain mutual exclusivity in statistical testing, each protein was assigned to one group without overlap between the groups. Conflicts in mutually exclusive classification were resolved by enforcing group priority from high functional specificity (e.g. nuclear pore) to low functional specificity (e.g. nucleus) in the following order: mitochondrial ribosome, yolk granule, lysosome, collagen, ribosome, nuclear pore, nuclear envelope, nucleolus, nucleoplasm, histone, nucleus, cytosol, mitochondrion, membrane, filament, filament and membrane, endoplasmic reticulum, extracellular, peroxisome, proteasome, translation, transcription, biosynthesis, and other. Group assignment was applied separately for each figure such that proteins were assigned only to the groups shown within the figure. Proteins with fold change z-score > 3 were excluded from analyses. A robust one-way ANOVA (Fig. 1e-f) or a robust two-way ANOVA (Fig. 2bc, 4d) (t1way and t2way from R package WRS2(v.1.1-4)) was performed to evaluate effects of genotype and protein group on the protein fold change during starvation (Table S1). Robust modeling was used to account for heteroscedasticity in proteomics data. Games-Howel post-hoc test was performed for proteomics group-wise comparisons (Table S1). For group-wise comparisons outside proteomics analyses, a within figure Benjamini-Hochberg multiple comparison corrected Wilcox two-sample t-test p-value is shown. A p-value < 0.05 or non-overlapping 95% confidence interval were considered statistically significant across all analyses.

## Acknowledgements

We thank Sebastian Leidel, Anat Bren, Peter Lenart, and Helge Grosshans for helpful feedback on the manuscript. We acknowledge support by the Microscopy Imaging Center at the University of Bern. Some strains were provided by the CGC, which is funded by NIH Office of Research Infrastructure Programs (P40 OD010440). Figures 1a-c and 3a were created with BioRender.com. Alleles *xe232, syb3768, syb1945, syb1880* and *syb1903* were created by SunnyBiotech.

## Author contributions

BDT and JT conceived the study, wrote the manuscript. JD, NS, BDT and JT edited the manuscript. JD and DSS conducted mass spectrometry measurements. JT and JP conducted all other experiments. JT and BDT performed computational and statistical analyses. BDT performed fitness modeling. SP implemented the U-net based image segmentation. NS designed alleles *syb1945, syb1880* and *syb1903*. JT, NS and JP created and validated strains used in the study.

## Data and Code availability

The mass spectrometry proteomics data have been deposited to the ProteomeXchange Consortium via the PRIDE^64^ partner repository with the dataset identifiers PXD050951 (label free), PXD067616 (TMT). Volume and fluorescence data from microscopy experiments is available in supplemental tables. Raw images are available upon request from the authors. Code for image analysis was as used in^24^ (10.5281/zenodo.10032867). Code for the population growth model is available as supplementary information.

## Supplementary Materials

Figures S1 to S6

Table S1

Data S1

