## Supplemental Information for "Autophagy-dependent proteome remodelling and ribosome decline balance starvation survival and recovery speed in *C. elegans*"

**Content**

Supplemental Figures S1 to S6
Supplemental Tables S1 to S2
Supplemental Data S1

**Fig. S1.**

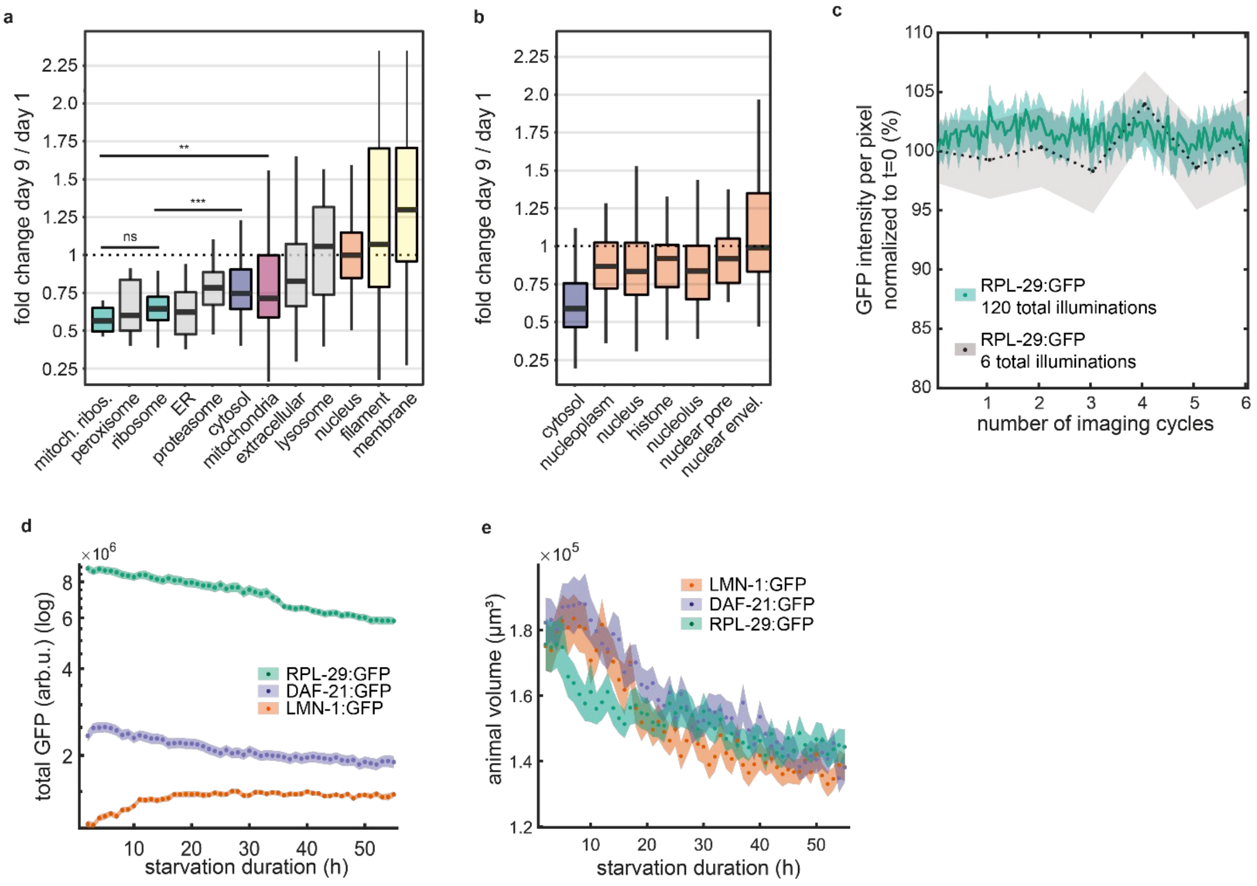


**Figure S1. a-b.** Box plots of rates of change measured for individual proteins in indicated groups for cellular organelles (**a**) and nuclear components (**b**) (quantified using LFQ). ***p<0.001 **p<0.01. Precise p-values and details on statistics in Supplemental Table S1 **c**. The effect of repeated imaging cycles on GFP signal strength. Test animals are illuminated 20 times more often than control animals using typical imaging settings (477 nm for 20ms at 30% power using Lumencor SpectraX light source). n = 116 test animals and 92 control animals. **d.** Total GFP-fluorescence of indicated strains during starvation. **e.** Animal volume during starvation. **c-e.** Mean with 95% CI (shaded area) **d,e.** n = 146, 323 and 137 animals from 3 to 4 biological repeats for LMN-1:GFP, DAF-21:GFP, and RPL-29:GFP.

### Fig. S2.

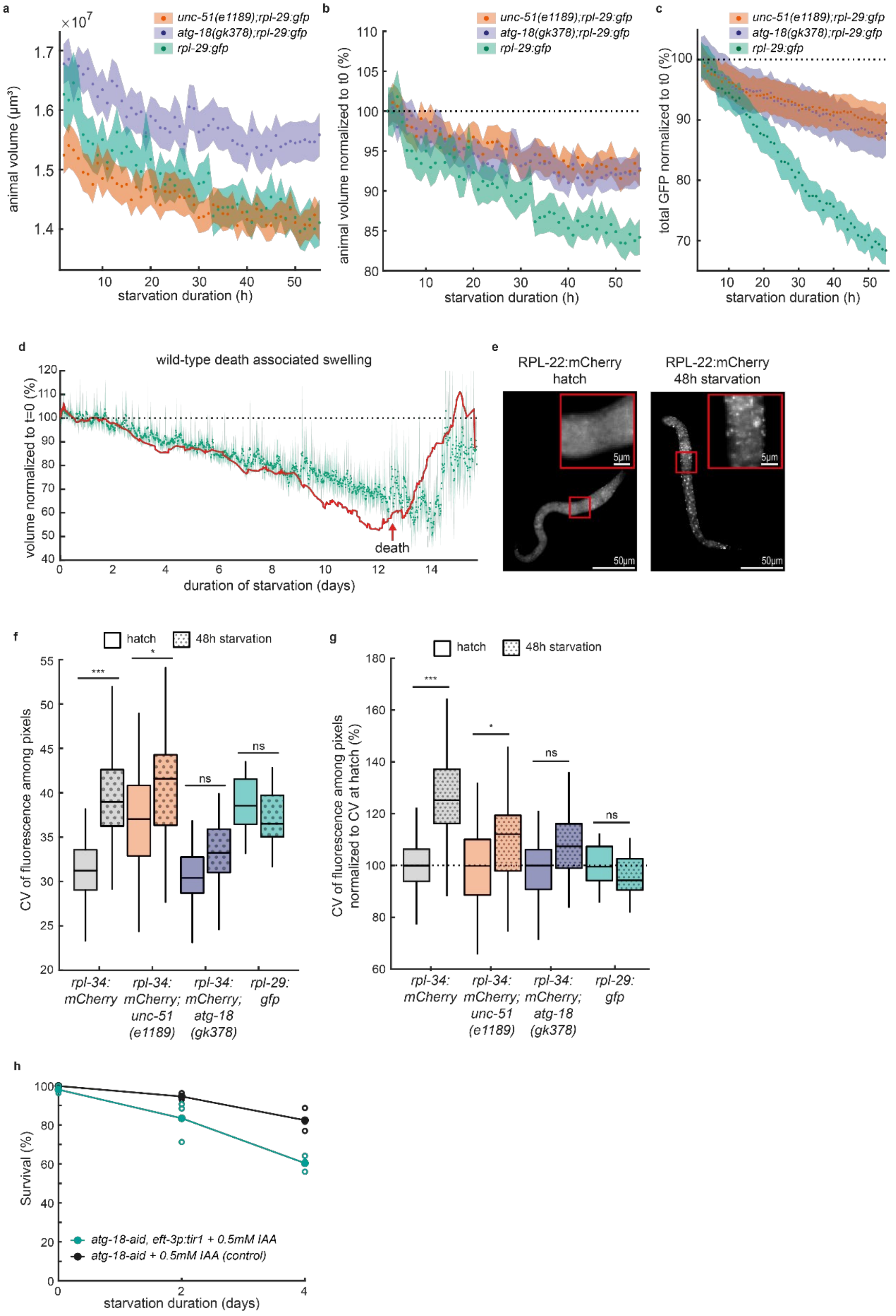

a
×10^7
unc-51(e1189);rpl-29:gfp
atg-18(gk378);rpl-29:gfp
rpl-29:gfp
animal volume (μm³)
starvation duration (h)
b
animal volume normalized to t0 (%)
starvation duration (h)
c
total GFP normalized to t0 (%)
starvation duration (h)
d
wild-type death associated swelling
volume normalized to t=0 (%)
death
duration of starvation (days)
e
RPL-22:mCherry
hatch
RPL-22:mCherry
48h starvation
5μm
50μm
f
hatch
48h starvation
CV of fluorescence among pixels
***
*
ns
rpl-34: mCherry
rpl-34: mCherry; unc-51 (e1189)
rpl-34: mCherry; atg-18 (gk378)
rpl-29: gfp
g
CV of fluorescence among pixels normalized to CV at hatch (%)
h
Survival (%)
atg-18-aid, eft-3p:tir1 + 0.5mM IAA
atg-18-aid + 0.5mM IAA (control)
starvation duration (days)

**Figure S2. a.** Volume of wild type and autophagy deficient mutants during starvation. **b.** Volume of wild type and autophagy deficient animals during starvation normalized to t=0. **c.** Total RPL-29:GFP fluorescence in wild type and in autophagy deficient animals during starvation. Mean and 95% CI of the mean for n = 283, 299, 101 animals from 3 biological repeats for *unc-51(e1189)* and *atg-18(gk378)* and 2 biological repeats for wild type **d**. Volume normalized to t=0 during starvation in wild-type animals and death associated swelling. Green: mean of 29 animals. Error bars: 95% CI. Representative individual animal is shown in red. Time of death is indicated by red arrow. **e**. RPL-22:mCherry in wild type at hatch and after 48h of starvation. **f.** Box plot of coefficient of variation (CV) among pixel intensities for strains and conditions shown in Fig. 2e (113, 49, 71, and 25 individuals per strain). Significance between group means (adjusted paired sample t-test): ***p<0.001, *p<0.05. Precise p-values in Table S2 **g.** Same as **f**, but normalized to t=0. **h.** Quantification of viability after ATG-18 AID. Animals were sampled from the same culture as used for mass spectrometry. Full circles: mean of n=3 biological repeats, empty circles: individual repeats. Statistics: p = 0.15; 0.15; 0.006 for 0, 2, and 4-day starvation (two-sample t-test) **a-c.** Mean with 95% CI (shaded area). n = 283, 299, 101 animals from 3 biological repeats for *unc-51(e1189)* and *atg-18(gk378)* and 2 biological repeats for wildtype.

**Fig. S3.**

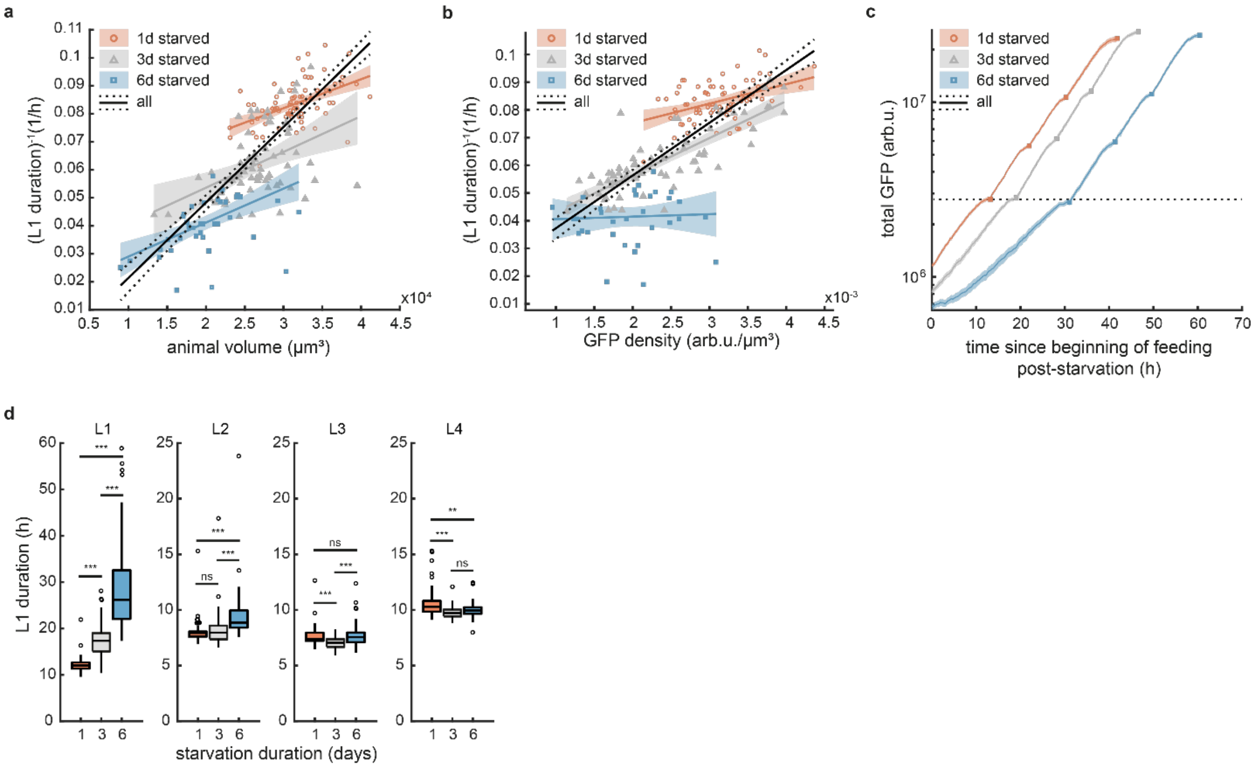


**Figure S3:** **a.** Animal volume vs. the inverse of the L1 larval stage duration after recovery from 1, 3 and 6-day starvations for 80, 70 and 37 individuals. Lines: linear regression with 95% CI as shaded area. Black: all starvation durations pooled ($r^2$=0.67). blue, gray, red: separated by starvation duration ($r^2$=0.28, $r^2$=0.17 and $r^2$=0.42 for 1, 3, and 6-days). **b**. GFP fluorescence volumetric density vs. the inverse of the L1 larval stage duration after recovery from 1, 3 and 6-day starvations for 79, 68 and 33 individuals. Lines: linear regression with 95% CI as shaded area. Black: all starvation durations pooled ($r^2$=0.64). blue, gray, red: separated by starvation duration ($r^2$=0.18, $r^2$=0.68 and $r^2$=0.01 for 1, 3, and 6-days). **c.** Mean GFP fluorescence after 1-, 3-, and 6-day starvation for 67, 66 and 43 individuals during post starvation feeding. Squares: molts between larval stages. Shaded area: 95% CI of the mean. Means were computed by aligning individual trajectories to the molts prioir to averaging, followed by re-scaling to the mean larval stage duration. Dotted line: mean L1 ribosome level after 1 day starvation. **d.** Box plots of all larval stage durations after 1, 3 , and 6 days of starvations (89, 89 and 71 individuals). Statistics: significance in difference between group means (adjusted two-sample t-test, ***p<0.001, *p<0.05). Precise p-values in Table S2.

**Fig. S4.**

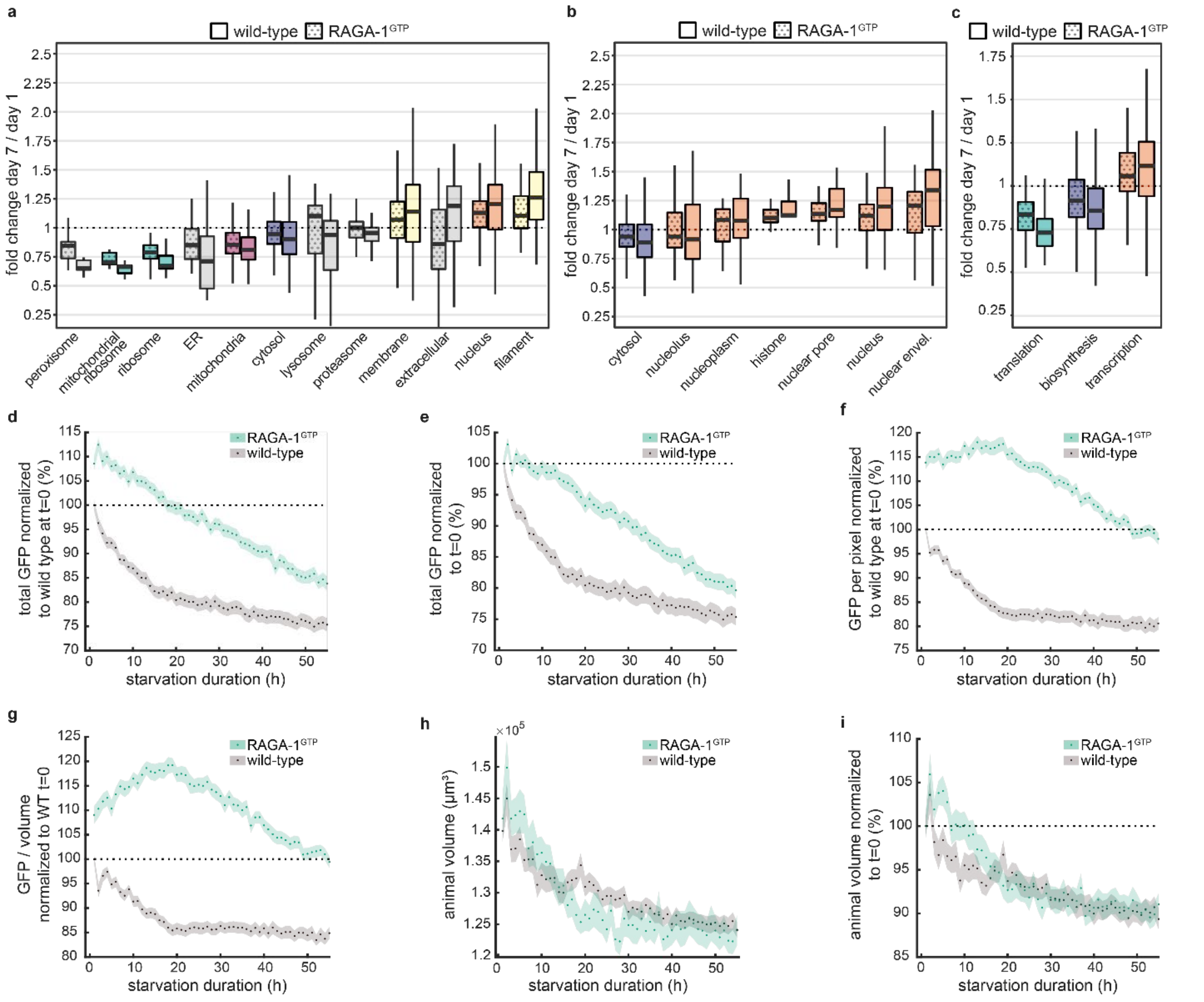


**Figure S4. a-c.** Box plots of fold change between day 1 and day 7 of startvation for individual proteins grouped by GO cellular component (**a,b**) and biological process (**c**) terms for wild-type (solid fill) and RAGA-1$^{GTP}$ (dotted fill). See Tables S1 for list of proteins in each group. **d**. Total GFP fluorescence normalized to wild-type t=0 during 55 hours of starvation in agarose chambers. **e**. Total GFP fluorescence normalized to t=0 of respective strain. **f**. Mean GFP fluorescence per pixel normalized to wild-type at t=0 **g**. GFP fluorescence volumetric density normalized to wild-type at t=0. **h**. Animal volume **i**. Animal volume normalized to t = 0. **d. – i.** n = 239 and 323 animals for wild type and RAGA-1$^{GTP}$ from 5 to 6 biological repeats. Circles are the mean among animals, shaded area is the 95% CI.

### Fig. S5.

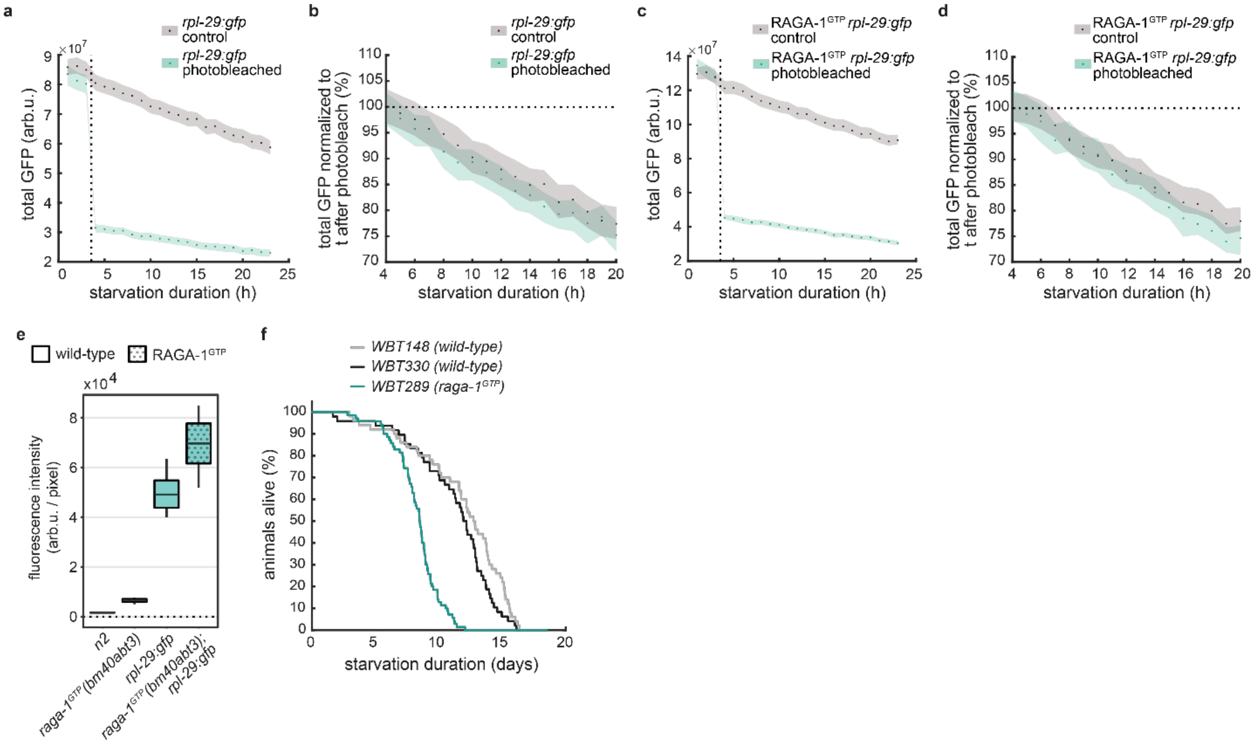


**Figure S5. a.** Effect of photo bleaching (477nm at full laser power for 8s) on total fluorescence in wild type animals. Dotted line is the timing of photobleaching. Control animals were not bleached. Circles are mean with 95% CI (shaded) n = 252 and 155 animals for bleached and control. **b.** same as a. but normalized to the first timepoint after photobleaching. Difference between rates of change in normalized fluorescence of control and photobleached animals would indicate GFP production, lack of difference shows absence of GFP production (see Methods). **c, d.** Same as a, b., but for in RAGA-1$^{GTP}$ animals. n = 113 animals for bleached and for control. **e**. Boxplot of N2 animal tissue autofluorescence, RAGA-1$^{GTP}$ associated GFP marker fluorescence, RPL-29:GFP fluorescence in wild type animals, and RPL-29:GFP fluorescence in RAGA-1$^{GTP}$ animals. n = 13 to 24 animals per condition. Animals were starved for 16 hours prior to immobilization and imaging on agarose pads. The background fluorescence from the agarose was substracted for all conditions. **f**. Starvation survival in agarose chambers in wildtype and RAGA-1$^{GTP}$ animals. The RAGA-1$^{GTP}$ strain WBT289 (see Methods) has an AID-tag associated with the RAGA-1$^{GTP}$ and expresses *tir-1(ubs38[F79G])* under an *eft-3* promoter. WBT148 does not express *tir-1* and has no AID-tag associated with *raga-1*. WBT330 contains the AID-tag on *raga-1* and expresses *tir-1(ubs38[F79G])* but is otherwise wild type for *raga-1*. Starvation survival is not affected by AID-tag or TIR-1 expression but is compromised by RAGA-1$^{GTP}$ mutation. 29 to 70 animals per strain.

**Fig. S6.**

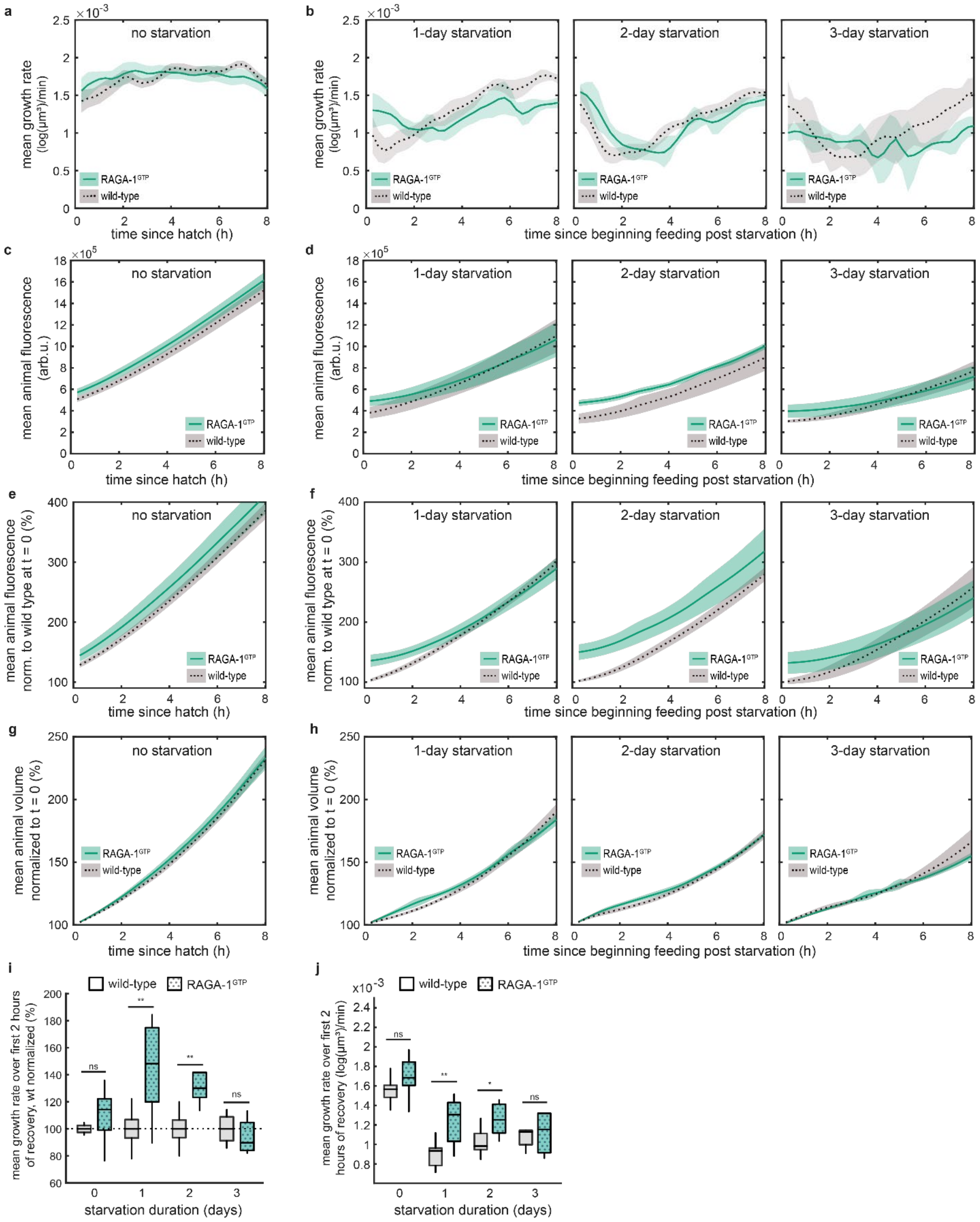


**Figure S6. a,b**. Relative growth rate ((dV/dt)/V) in RAGA-1$^{GTP}$ and wild type animals during *ad libitum* feeding after hatching (a) and after indicated duration of starvation (b) for the first 8 hours of feeding. Mean and 95% CI (shaded area). **c,d**. As a,b but for total fluorescence **e,f**. As a, b but for total fluoresccence normalized to wild type fluorescence at t=0 **g,h**. as a,b but for volume **i**. Box plots of the ratio of relative growth rate (($\Delta V/\Delta t$)/V) between RAGA-1$^{GTP}$ and wild type (or between wild type and mean of wild type) animals during *ad libitum* feeding after hatching and after indicated duration of starvation over the first 2 hours of feeding. Statistics: comparison between group means (adjusted paired sample t-test, $**p<0.01$, $*p<0.05$). Precise p-values in Table S2 **j**. as i. but without normalization to wild type. Data from 177 to 386 individuals across 2 to 5 biological repeats per strain and condition.

**Table S1.**

Precise p-values and details of statistical test applied to proteomics experiments (all figures). See separate file.

**Table S2.**

Precise p-values and details of statistical test applied to experiments not involving proteomics (all figures). See separate file.

### Data S1.

```
deltas = linspace(0.01, 0.15, 20);%increase in lag per starvation (unitless, e.g. in hours  per hour)
t_starvations = linspace(0,20,21)*24; % starvation durations in hours

k = 3500;


Tstart = 40; % start of reproduction
rho = 6; %eggs per hour
n=4; % Hill functionof death curve

bottleneck = 20*numel(deltas); %total capacity of food: 20*starting population. end of simulation

for t_starv_ind = 1:numel(t_starvations) %loop over different starvatoin durations
    t_starvation = t_starvations(t_starv_ind);
    disp(t_starv_ind);
    for delta_ind = 1:numel(deltas) %loop over different survival times
        delta = deltas(delta_ind);
        for t = 1:1000 % run for 1000 hours
           rep(t,delta_ind,t_starv_ind) = reproduction(Tstart, t, t_starvation, delta, k, rho, n); %compute reproduction in the last hour
           totrep(t,delta_ind,t_starv_ind) = sum(rep(1:t,delta_ind,t_starv_ind)); % comput total reproduction so far per "tradeoff strategy"
        end
    end
end

tot_tot_rep = squeeze(sum(totrep,2)); %total reproduction across all tradeoff-strategies

for t_starv_ind = 1:numel(t_starvations)
    t_lim(t_starv_ind) = min(find(tot_tot_rep(:,t_starv_ind) >bottleneck)); % for each starvation duration find time-point when food is
exhausted
end

for delta_ind = 1:numel(deltas)
    for t_starv_ind = 1:numel(t_starvations)
        totrepattlim = totrep(t_lim(t_starv_ind),delta_ind,t_starv_ind); %caluculate total reproduction when food is exhausted per strategy
        tot_tot_rep_at_tlim = tot_tot_rep(t_lim(t_starv_ind),t_starv_ind) %calculate total number of worms when food is exhausted
        fitness(delta_ind, t_starv_ind) = totrepattlim/tot_tot_rep_at_tlim; %frequency of strategy at the time of food exhaustion
    end
end
med_survival=deltas*k/24; %median survival times for a given strategy in days
lagtimes = deltas*24 %lag time increase hours per day of starvation

%%
figure
imshow(fitness,[0,0.15])
colorbar

%%
function tau = lag_time(t,delta) %calculation of lag time for a given survival rate and proportionality factor
        tau = t*delta;
end

function s = survival(t,delta,n,k) %survival rate
        s = 1/(1+(t/(delta*k))^n);
end

function r_t = reproduction(Tstart, t_recovery, t_starvation, delta, k, rho, n) % reproduction, taking into account survival rate
        if t_recovery>(Tstart+lag_time(t_starvation,delta)) & t_recovery <  (Tstart+lag_time(t_starvation,delta) + 48) %test if lag already
over and if reproductive phase already over
            r_t = survival(t_starvation,delta, n, k)*rho; %compute reproduction by fraction number of survivors * eggs/hour
        else
            r_t = 0;
        end
end
```

Data S1: Matlab code for the population growth model presented in Fig. 4i.
